# Human iPSC-derived myelinating organoids and globoid cells to study Krabbe Disease

**DOI:** 10.1101/2024.07.19.604372

**Authors:** Lisa Marie P Evans, Joseph Gawron, Fraser J Sim, M Laura Feltri, Leandro N Marziali

**Author notes:** Deceased (M Laura Feltri). Correspondence to: Leandro N Marziali Full address: 701 Ellicott St., Buffalo, NY, 14203, USA.

## Abstract

Krabbe disease (Kd) is a lysosomal storage disorder (LSD) caused by the deficiency of the lysosomal galactosylceramidase (GALC) which cleaves the myelin enriched lipid galactosylceramide (GalCer). Accumulated GalCer is catabolized into the cytotoxic lipid psychosine that causes myelinating cells death and demyelination which recruits microglia/macrophages that fail to digest myelin debris and become globoid cells. Here, to understand the pathological mechanisms of Kd, we used induced pluripotent stem cells (iPSCs) from Kd patients to produce myelinating organoids and microglia. We show that Kd organoids have no obvious defects in neurogenesis, astrogenesis, and oligodendrogenesis but manifest early myelination defects. Specifically, Kd organoids showed shorter but a similar number of myelin internodes than Controls at the peak of myelination and a reduced number and shorter internodes at a later time point. Interestingly, myelin is affected in the absence of autophagy and mTOR pathway dysregulation, suggesting lack of lysosomal dysfunction which makes this organoid model a very valuable tool to study the early events that drive demyelination in Kd. Kd iPSC-derived microglia show a marginal rate of globoid cell formation under normal culture conditions that is drastically increased upon GalCer feeding. Under normal culture conditions, Kd microglia show a minor LAMP1 content decrease and a slight increase in the autophagy protein LC3B. Upon GalCer feeding, Kd cells show accumulation of autophagy proteins and strong LAMP1 reduction that at a later time point are reverted showing the compensatory capabilities of globoid cells. Altogether, this supports the value of our cultures as tools to study the mechanisms that drive globoid cell formation and the compensatory mechanism in play to overcome GalCer accumulation in Kd.

## Introduction

Krabbe Disease (Kd), also known as globoid cell leukodystrophy, is a lysosomal storage disease (LSD) cause by mutations in the *GALC* gene which encodes the lysosomal hydrolase galactosyl ceramidase (GALC). The incidence of Kd in USA is estimated to be 1:100,000-1:250,000 (Barczykowski et al., 2012). Kd is classified on the basis of the age of symptoms onset as early infantile, late infantile, juvenile and adult (Bascou et al., 2018; Duffner et al., 2011; Duffner et al., 2012). Kd patients experience rapid and severe demyelination, neurodegeneration, and neuroinflammation in the central nervous system (CNS) and peripheral nervous system (PNS). Symptoms include: progressive neurologic deterioration with decorticate posturing, blindness, deafness, peripheral neuropathy, autonomic instability, and death (Duffner et al., 2011; Duffner et al., 2012).

White matter is mostly composed of axons that are ensheathed by myelin which is produced by mature oligodendrocytes (OLs) in the CNS. Myelin is a lipid-rich insulating multi-layer that protects and provides trophic support to axons, and allows rapid impulse propagation through the saltatory propagation of action potentials between Nodes of Ranvier (Simons et al., 2024). Oligodendrocyte Progenitor Cells (OPCs) arise from neuroectoderm derived Neural Stem Cells (NSCs) which also produce neurons and astrocytes (El Waly et al., 2014). Microglia, the innate immune system surveillants of the CNS, originate from the mesoderm derived yolk sac which produces microglial progenitors capable of migrating towards and colonizing the CNS during embryonic development (Sierra et al., 2024).

Animal models of Kd are available and in particular, the twitcher mouse (Kobayashi et al., 1980), a global *GALC* knock out model, and the recently developed *GALC* conditional knock out mouse (Weinstock, Shin, et al., 2020) together with human biopsies had been used to study the physiopathology of Kd. In Kd, the lack of GalC function or the failure to dock GalC into lysosomes (Shin et al., 2016) causes the accumulation of GalC’s substrate galactosyl ceramide (GalCer) which is alternatively deacylated by the catabolic enzyme acid ceramidase (Asah1) to produce psychosine (Li et al., 2019), a very potent toxin. The myelinating cells, OLs in the CNS and Schwann Cells (SC) in the PNS, produce large amounts of GalCer and are particularly susceptible to cell death due to the accumulated psychosine. The subsequent demyelination triggers an inflammatory response that recruits microglia/macrophages (CNS/PNS) to demyelinating foci. Microglia/macrophages phagocytize and digest GalCer rich myelin debris however, the lack of GalC causes GalCer and psychosine accumulation, and the formation of multinucleated cells named globoid cells.

Whereas animal models of Kd give us the opportunity to study the dynamics of the cellular and molecular events that drive the disease and to test different therapeutic interventions, the study of the human disease is limited to post mortem tissue. In the past few years, the utilization of iPSCs to replicate organogenesis *in vitro* has emerged and several diseases have been successfully modeled providing the tools to perform in-detail studies of the cellular and molecular mechanisms that drive different diseases (Adlakha, 2023; Eichmuller & Knoblich, 2022). Here, we show that Kd can be modeled by using myelinating organoids that show no obvious defects in neurogenesis, astrogenesis and oligodendrogenesis but signs of demyelination and iPSC-derived microglia that when fed with GalCer give rise to globoid cells.

## Materials and methods

### iPSC demographics and culture

All iPSC lines belong to de-identified donors and all the procedures performed in the present study were approved by IRB (MOD00014209). We used 4 iPSC lines including 2 Control (Ctrol) lines from apparently healthy individuals and 2 iPSC lines from early onset Kd patients. The 2 Ctrol and 1 of the Kd lines were purchased from The Coriell Institute: GM23720 iPSC from B-lymphocyte of an apparently healthy 22-year-old female, AG27875 iPSC from fibroblast of an apparently healthy 29-year-old male and GM26644 iPSC from fibroblast of a two-year-old female Kd patient homozygous for the previously characterized 30kb deletion of the *GALC* gene (Luzi et al., 1995). The second Kd line was a gift from Dr. Daesung Shin at SUNY at Buffalo: this line was an iPSC from fibroblast produced at The Induced Pluripotent Stem Cell Generation facility at SUNY at Buffalo; the originating fibroblasts were obtained from the Telethon Biobank (FFF0751981) with unknown demographics bearing compound heterozygous *GALC* mutations (30kb-del + T529M) (Rafi et al., 1995; Shin et al., 2016).

iPSCs were cultured on non-cell culture treated plates (Greiner Bio-One 657185) coated with 8.6 µg/cm^2^ Growth Factor Reduced Matrigel (Corning 356230) and mTeSR-Plus media (STEMCELL Technologies, 05825) at 37°C/5%CO_2_. Upon thawing iPSCs were plated in media containing 10 µM Y27632 - ROCK inhibitor (STEMCELL Technologies, 72304) for 24 hrs. Afterwards, ROCK inhibitor was removed from the media and iPSCs passed when reached 80% confluency using ReLeSR Reagent (STEMCELL Technologies, 05872).

### Myelinating organoid culture

Spinal cord patterned myelinating organoids were generated as described previously (James et al., 2021). iPSC colonies were detached with 15 U/ml Dispase II (Thermo Fischer Scientific, 17105-041) in PBS for 10-15 min in the incubator. After washing with PBS, colonies were transferred into Ultra-Low Attachment 6 well plates (Corning, 3473) in media containing Phase I media: 0.5X Iscove’s Modified Dulbecco’s Medium (Thermo Fischer Scientific, 12440053), 0.5X Ham’s F-12 Nutrient Mixture (Thermo Fischer Scientific, 11765054), 0.5% BSA, 1X Chemically Defined Lipid Concentrate (Thermo Fischer Scientific, 11905031), 0.004% (v/v) monothioglycerol (Sigma, M6145), 7 µg/ml insulin (Sigma, 11376497001), 15 µg/ml transferrin (Sigma, 10652202001), 1X Antibiotic-Antimycotic (Sigma-Aldrich, A5955), 1 mM N-acetyl cysteine (Sigma, A8199), 10 µM SB431542 (STEMCELL Technologies, 72234), 0.1 μM LDN-193189 (STEMCELL Technologies, 72147) and cultured at 37°C/5%CO_2_. On day 7, spheroids were moved to Phase II media: Phase I media without SB431542 and LDN-193189, and the addition of 0.1 µM retinoic acid (RA, Sigma, R2625), 5 ng/ml FGF-2 (PeproTech, AF-100-18B) and 5 ng/ml heparin (Sigma, H3149), and cultured at 37°C/5%CO_2_. On day 14, spheroids were plated onto 25 µg/mL poly-L-lysine/10 µg/mL laminin (Sigma, P5899 and L2020) coated plates in Phase II media and cultured at 37°C/5%CO_2_. On day 17, spheroids were lifted and returned to suspension culture in Phase III media: 1X Advanced DMEM/F12 (Thermo Fisher Scientific, 12491015), 0.5X N2 (Thermo Fisher Scientific, 1750200), 0.5X B27 (Thermo Fisher Scientific, 17504-044), 0.5X Glutamax (Thermo Fisher Scientific, 35050061), 1X Antibiotic-Antimycotic, 5 ng/ml FGF-2, 5 ng/ml heparin, 1 µM RA, 1 µM purmorphamine (STEMCELL Technologies, 72204), and cultured at 37°C/10%CO_2_. After 8 days, FGF-2 and heparin were removed from the media. After 14 days, Media was changed to oligodendrocyte proliferation media: 1X Advanced DMEM/F12, 1X N2, 0.5X B27, 1X Glutamax, 1X Antibiotic-Antimycotic, 10 ng/ml FGF-2, 20 ng/ml PDGF-AA (PeproTech, AF-100-13A), 1 µM purmorphamine, 1 µM smoothened agonist (EMD Millipore, 566660), 10 ng/ml IGF-1 (PeproTech, AF-100-11), 45 ng/ml T3 (Sigma, T6397), 5 µg/ml heparin and cultured at 37°C/7.5%CO_2_. After 3 weeks, spheroids were transferred to a PTFE-coated Millicell Cell Culture Inserts in 6 well plates, and maintained in myelination media: 1X Advanced DMEM/F12, 1X N2, 0.5X B27, 1X Glutamax, 1X Antibiotic-Antimycotic, 1X ITS-X (Thermo Fisher Scientific, 51500056), 10 ng/ml IGF-1, 45 ng/ml T3 and 5 µg/ml heparin and cultured at 37°C/7.5%CO_2_. Media was changed every 2-3 days throughout the protocol.

### Microglia Culture

iPSC-derived microglia were produced as described previously (Douvaras et al., 2017). iPSCs were dissociated to single cell suspension with Accutase^®^ (Sigma, A6964) for 2-3 min. in the incubator and plated onto 8.6 µg/cm^2^ Matrigel (Corning, 356234) coated dishes at a density of 1.5×10^3^ cells/cm^2^ in mTeSR-Plus media containing 10μM Rock Inhibitor for 24 hrs. After 1-2 days, when colonies became visible, media was changed to mTeSR Custom media (STEMCELL Technologies), containing 80 ng/ml BMP4 and cultured at 37°C/5%CO_2_ with daily media changes for 4 days. Next, media was changed to StemPro-34 SFM, 1X GlutaMAX-I, 25 ng/ml FGF-2, 100 ng/ml SCF (PeproTech, 300-07) and 80 ng/ml VEGF (PeproTech, 100-20). After 2 days, media was changed to StemPro-34, 50 ng/ml SCF, 50 ng/ml IL-3 (PeproTech, 200-03), 5 ng/ml TPO (PeproTech, 300-18), 50 ng/ml M-CSF (PeproTech, 300-25) and 50 ng/ml Flt3 ligand (R&D Systems, 308-FK-100/CF). From this point onwards, cells in the supernatant were pelleted and returned to the culture with fresh media every 3-4 days. On day 14, media was changed to StemPro- 34, 50 ng/ml M-CSF, 50 ng/ml Flt3 ligand and 25 ng/ml GM-CSF (PeproTech, AF-300-03). From day 25 to 45, microglia progenitors were isolated by Magnetic Cell Sorting (MACS) with CD14 beads (Miltenyi Biotec, 130-118-906), plated onto tissue culture-treated dishes or Thermanox plastic coverslips in media containing RPMI-1640, 1X GlutaMAX-I, 10 ng/ml GM-CSF and 100 ng/ml IL-34 (PeproTech, 200-34), and cultured at 37°C/5%CO_2_ with media changes every 2-3 days for at least 10 days.

### EdU incorporation

Proliferating cells were labelled with the thymidine analogue 5-ethynyl-2′-deoxyuridine (EdU) (Sigma; 900584) at a 1 µM concentration that was added to the culture media 1 h before fixation. EdU incorporation was visualized by using click chemistry. Briefly, fixed cells were incubated for 30 min at RT with a solution containing 8 µM Alexa Fluor 488-azide (Cedarlane; CLK-1275-1), 2 mM CuSO_4_·5H_2_O, and 20 mg/ml ascorbic acid.

### GalC Activity

GalC activity was measured as previously described (Martino et al., 2009). Briefly, proteins were extracted using a solution containing 10 mmol/L Sodium Phosphate buffer pH 6.0 and 0.1% (v/v) Nonidet NP-40 for 30 min on ice followed 3 rounds of sonication and a 10 min. centrifugation at 13,000g. Next, supernatants were collected and protein concentration measured by BCA assay (Thermo Fischer Scientific). GalC activity assays were run using 5 µg of protein (50 µL) that were added to a substrate mixture containing: 100 µL of 1.5 mmol/L 4-Methylumbelliferyl-β-D-galactopyranoside (MUGAL) substrate diluted in 0.1/0.2 mol/L citrate/phosphate buffer pH 4.0 and 15 μM AgNO_3_. The reactions were run for 30 minutes at 37°C and stopped by adding 100 µL of 0.2 mol/L Glycine/NaOH, pH 10.6. Next, fluorescence was measured using a BioTek Cytation 5 spectrofluorometer (λ_ex_ 360 nm, λ_em_ 446 nm) and MUGAL break down was determined by quantifying the amount of 4-Methylumbelliferone liberated utilizing a 4-Methylumbelliferone (Sigma, M1381) standard curve. Enzymatic activity was calculated as 1 unit been 1.0 μmol/min of substrate hydrolyzed at 37°C.

### Immunohistochemistry and Immunocytochemistry

For immunocytochemistry, organoids were fixed with 2% paraformaldehyde (PFA) in PBS at 4°C ON. Next, the organoids were rinsed with PBS, cryopreserved by immersion in 30% sucrose until sunk and cryosectioned with a Leica CM1850 cryostat to obtain 20 μm thick sections. Sections were blocked for 2 hrs. at RT with 5% fetal bovine serum (FBS) in PBS containing 0.2% Triton X-100. Primary antibodies (See Table 1) were incubated ON at 4°C followed by 3 washes with PBS and the corresponding secondary antibodies (1:500; Jackson ImmunoResearch Laboratories) containing 4′,6-diamidino-2-phenylindol (DAPI; 1 μg/ml) were incubated for 2 hrs. at RT. After 3 washes with PBS, sections were mounted with Epredia™ Immu-Mount™ (Fisher Scientific) and imaged with a Zeiss Apotome microscope and analyzed using ImageJ software.

**Table 1.**
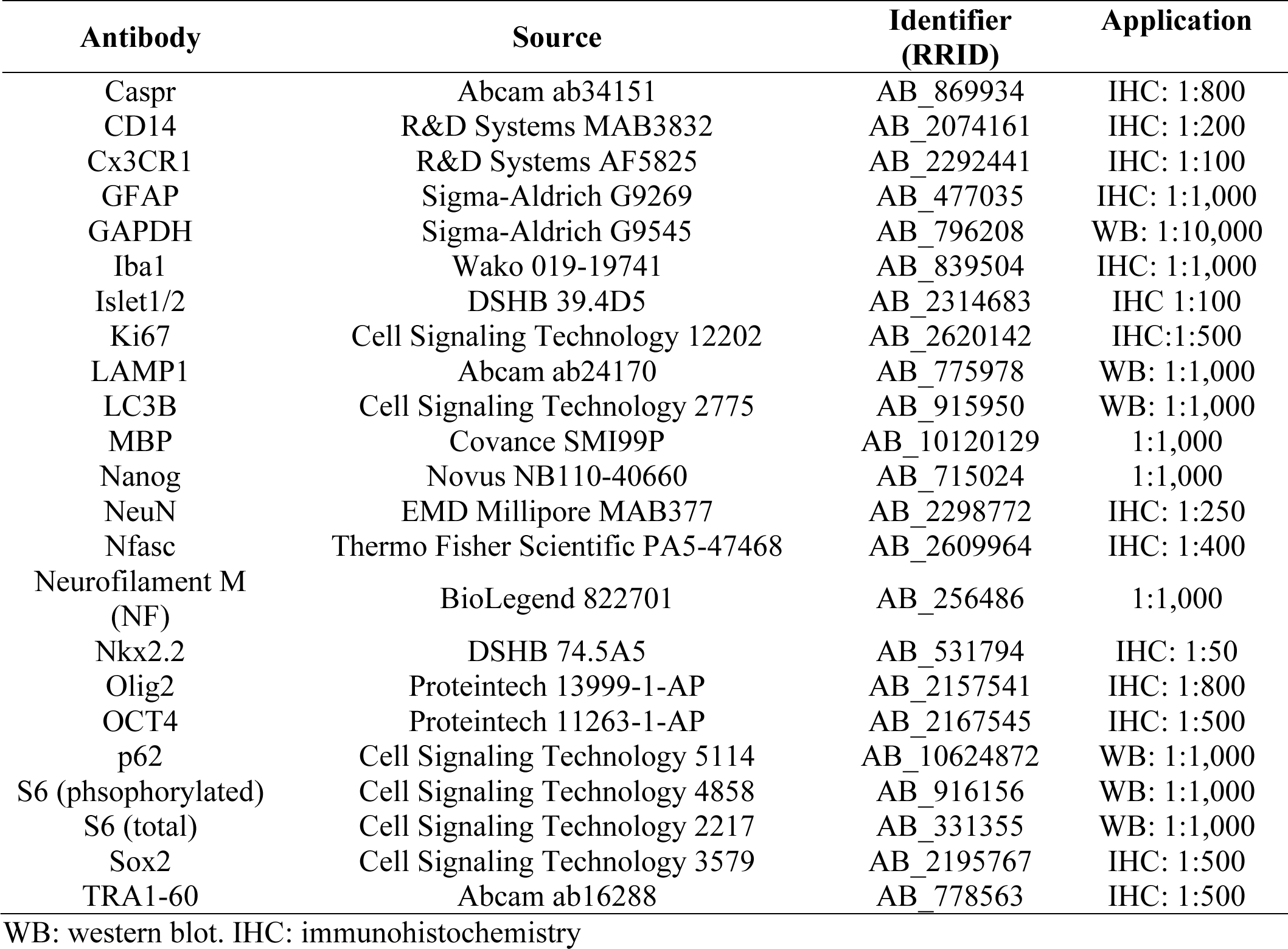
Antibodies.

| **Antibody** | **Source** | **Identifier (RRID)** | **Application** |
| --- | --- | --- | --- |
| Caspr | Abcam ab34151 | AB_869934 | IHC: 1:800 |
| CD14 | R&D Systems MAB3832 | AB_2074161 | IHC: 1:200 |
| Cx3CR1 | R&D Systems AF5825 | AB_2292441 | IHC: 1:100 |
| GFAP | Sigma-Aldrich G9269 | AB_477035 | IHC: 1:1,000 |
| GAPDH | Sigma-Aldrich G9545 | AB_796208 | WB: 1:10,000 |
| Iba1 | Wako 019-19741 | AB_839504 | IHC: 1:1,000 |
| Islet1/2 | DSHB 39.4D5 | AB_2314683 | IHC 1:100 |
| Ki67 | Cell Signaling Technology 12202 | AB_2620142 | IHC:1:500 |
| LAMP1 | Abcam ab24170 | AB_775978 | WB: 1:1,000 |
| LC3B | Cell Signaling Technology 2775 | AB_915950 | WB: 1:1,000 |
| MBP | Covance SMI99P | AB_10120129 | 1:1,000 |
| Nanog | Novus NB110-40660 | AB_715024 | 1:1,000 |
| NeuN | EMD Millipore MAB377 | AB_2298772 | IHC: 1:250 |
| Nfasc | Thermo Fisher Scientific PA5-47468 | AB_2609964 | IHC: 1:400 |
| Neurofilament M (NF) | BioLegend 822701 | AB_256486 | 1:1,000 |
| Nkx2.2 | DSHB 74.5A5 | AB_531794 | IHC: 1:50 |
| Olig2 | Proteintech 13999-1-AP | AB_2157541 | IHC: 1:800 |
| OCT4 | Proteintech 11263-1-AP | AB_2167545 | IHC: 1:500 |
| p62 | Cell Signaling Technology 5114 | AB_10624872 | WB: 1:1,000 |
| S6 (phosphorylated) | Cell Signaling Technology 4858 | AB_916156 | WB: 1:1,000 |
| S6 (total) | Cell Signaling Technology 2217 | AB_331355 | WB: 1:1,000 |
| Sox2 | Cell Signaling Technology 3579 | AB_2195767 | IHC: 1:500 |
| TRA1-60 | Abcam ab16288 | AB_778563 | IHC: 1:500 |

WB: western blot. IHC: immunohistochemistry

For myelin labelling, non-sectioned whole organoids were stained. After fixation and cryopreservation, organoids were blocked with 5% FBS in PBS containing 1% Triton X-100 for 24 hrs. at 4°C. Primary antibodies (See Table 1) were diluted in 2% FBS in PBS 1% Triton X-100 and incubated for 24 hrs. at 4°C followed by 4 washes (30 min each) with PBS. The corresponding secondary antibodies (1:500; Jackson ImmunoResearch Laboratories) were diluted in in 2% FBS in PBS 1% Triton X-100 containing DAPI (1 μg/ml) and incubated for 24 hrs. at 4°C. After 4 washes (30 min each) with PBS, organoids were individually place in 96 well glass bottomed plates (Corning, 4580) containing PBS and imaged using a Leica DMI6000 CS confocal microscope. Stacks were processed using ImageJ’s plug in Simple Neurite Tracer’s (SNT) to count the number and measure length of myelin internodes (Arshadi et al., 2021).

### Immunoblotting

Total proteins were extracted using RIPA buffer supplemented with protease and phosphatase inhibitors. Proteins (1–2 μg) were loaded into Nupage™ 4-12% Bis Tris Protein Gels, 1.0 mm, 17 wells or in house made 4-12% Bis-Tris, 1.0 mm, 21 wells and run using MES buffer. Transfer onto polyvinylidene difluoride (PVDF) membranes was done using BioRad’s Trans-Blot Turbo Transfer System. The membranes were blocked with 5% Bovine Serum Albumin (BSA) in TBS containing 0.2% Tween 20 for 2 hrs. at RT. Primary antibodies (See Table 1) were diluted in blocking solution and incubated ON at 4°C. The corresponding peroxidase-conjugated secondary antibodies were diluted in blocking solution and incubated for 2 hrs. at RT. The blots were developed by using ECL Select Western Blotting Detection Reagent (Cytiva’s Amersham) and a ChemiDoc XRS+ system (Bio-Rad).

### GalCer feeding

Mature microglia were exposed to 10 μM GalCer (C8 Galactosyl(β) Ceramide (d18:1/8:0)) (Avanti Polar Lipids, 860538P) for 16, 24 or 48 hrs. GalCer was resuspended at 20 mM concentration in DMSO, stored in a N_2_ inert atmosphere and used no more than 2 times. To facilitate GalCer incorporation, the lipid carrier (2-Hydroxypropyl)-beta-cyclodextrin (Sigma, 778966) was used at a 0.1 mM final concentration.

### Electron microscopy

Organoids were fixed with 2.5% glutaraldehyde/4% PFA in 0.12 M phosphate buffer (pH 7.4) for at least 24 hrs. at 4°C. Next, the samples were stained with 1% osmium tetroxide for 2 hrs. at RT, dehydrated, and embedded into epoxy resin (Sigma-Aldrich; 45359-1EA-F). Ultrathin (90 nm) sections were obtained with a Leica EM UC7 ultramicrotome. Ultrathin sections were stained with uranyl acetate and lead citrate and imaged with an FEI Tecnai G2 Spirit BioTwin transmission electron microscope.

### Statistical analysis

Measurements belonging to individual Ctrol or Kd lines were grouped and colour coded to reflect that each value represents a replicate of an individual culture/organoid. Each experiment was repeated at least 2 times. GraphPad Prism 9.1.1 software was used to perform two-tailed Student’s *t* tests to compare two groups/time points and two-way ANOVAs followed by multiple-comparisons post-tests to compare more than two groups and time points. Statistical significance was set at a *P* value of <0.05.

## Results

### Kd iPSCs do not show any morphological and proliferation defects

To model Kd in vitro for the study of the pathological mechanisms of Kd, we used iPSCs derived from Kd dermal fibroblasts. We first confirmed that Kd iPSCs have a significant reduction in GalC activity when compared to Ctrol ones (**Figure 1A**). Next, we observed that Kd iPSCs form normally appearing compact colonies with well-defined borders, no morphological signs of differentiation and express the pluripotency factors Nanog, TRA1-60 and OCT4 similarly to Ctrol (**Figure 1B**). Finally, Kd iPSCs show no difference in their proliferation rate when compared to Ctrol ones as shown by EdU pulse (**Figure 1C**).

**Figure 1.**
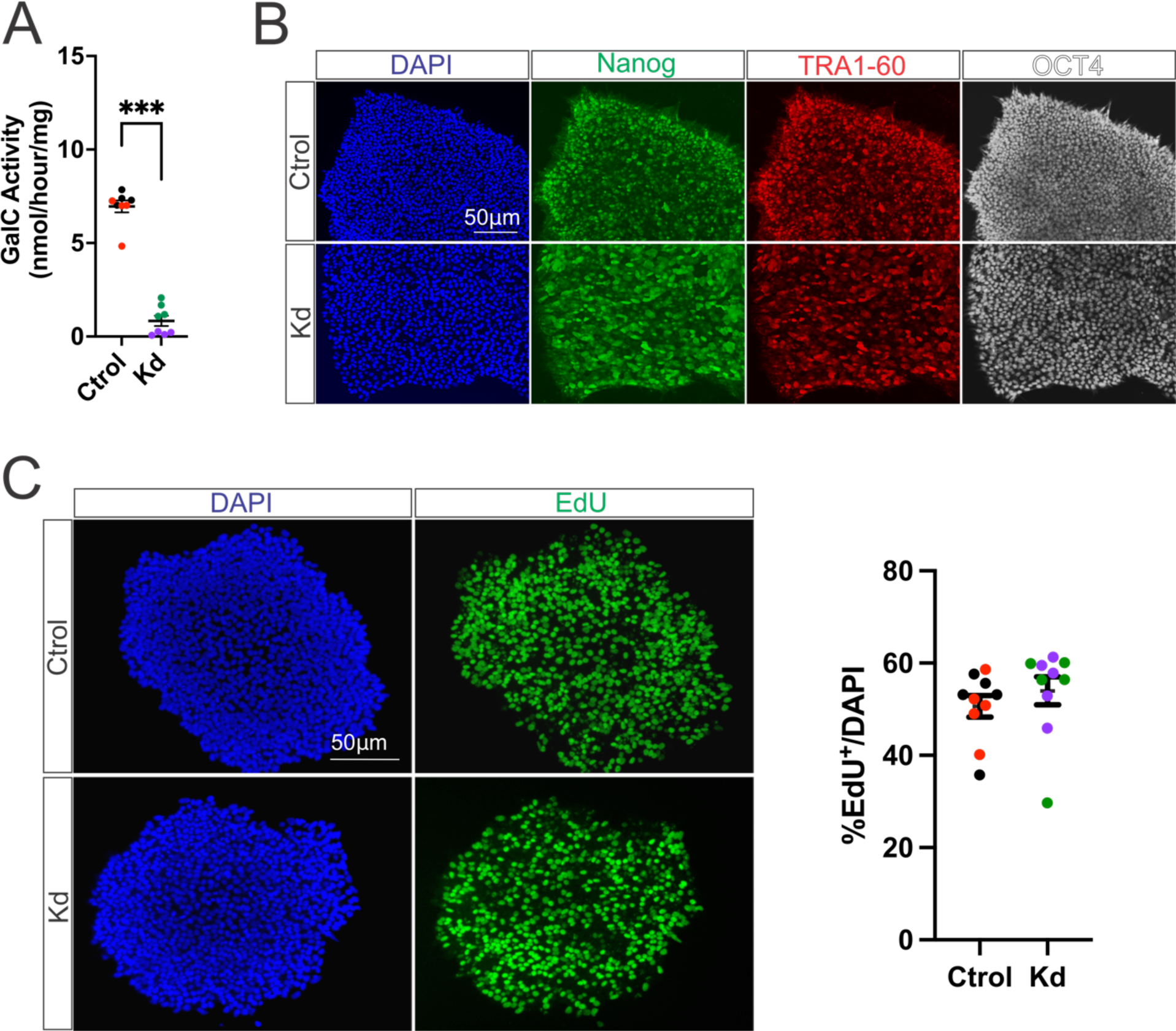
Kd iPSCs do not show any morphological and proliferation defects. (A) GALC activity measurements of Ctrol and Kd iPSCs. (B) Immunohistochemistry of Ctrol and Kd iPSCs for the pluripotency markers Nanog, TRA1-60 and OCT4. (C) Proliferation assessment by EdU incorporation assay of Ctrol and Kd iPSCs. Colors indicate replicates from individual iPSC lines. Values are expressed as the means ± SEMs. ***P < 0.001 by Student’s t test.

### Kd organoids develop normally and show no defects in neurogenesis, astrogenesis, and oligodendrogenesis

To study neurodevelopment and myelination in Kd, we used a previously described protocol to make iPSC-derived myelinating organoids that recapitulate spinal cord development (**Figure 2A**) (James et al., 2021). Under these culture conditions, iPSCs first give rise to neuroectoderm (D7-20) followed by neurogenesis (D37), astrogenesis (D37), oligodendrogenesis (D37-60) and later on, myelination (myelination weeks 0-28 counting from D60). At a glance, Kd organoids show no macroscopic differences after 60 days in culture when compared to Ctrol (**Figure 2B**). Next, we analyzed the NSC population by Sox2 staining and found no differences between Ctrol and Kd organoids at D20 (**Figure 2C**). At D37 there is no difference in the number of proliferating Ki67^+^ cells (**Figure 2D**) supporting the idea that Kd organoids develop similarly to Ctrol. In regards to neurogenesis, we found no differences between groups in the number of NeuN^+^ neurons and Islet-1/2^+^ neuronal progenitors at D37 (**Figure 2E and 2F**). Astrogenesis was similar between groups as shown by GFAP content at D37 (**Figure 2G**). In regards to oligodendrogenesis, we analyzed 2 time points: D37 and D60, the first one representing endogenous oligodendrogenesis and the second one oligodendrogenesis driven by the exposure of the cultures to the oligodendrocyte specific growth factors PDGF-AA and FGF-2. In both cases, we found no differences between groups (**Figure 2H**).

**Figure 2.**
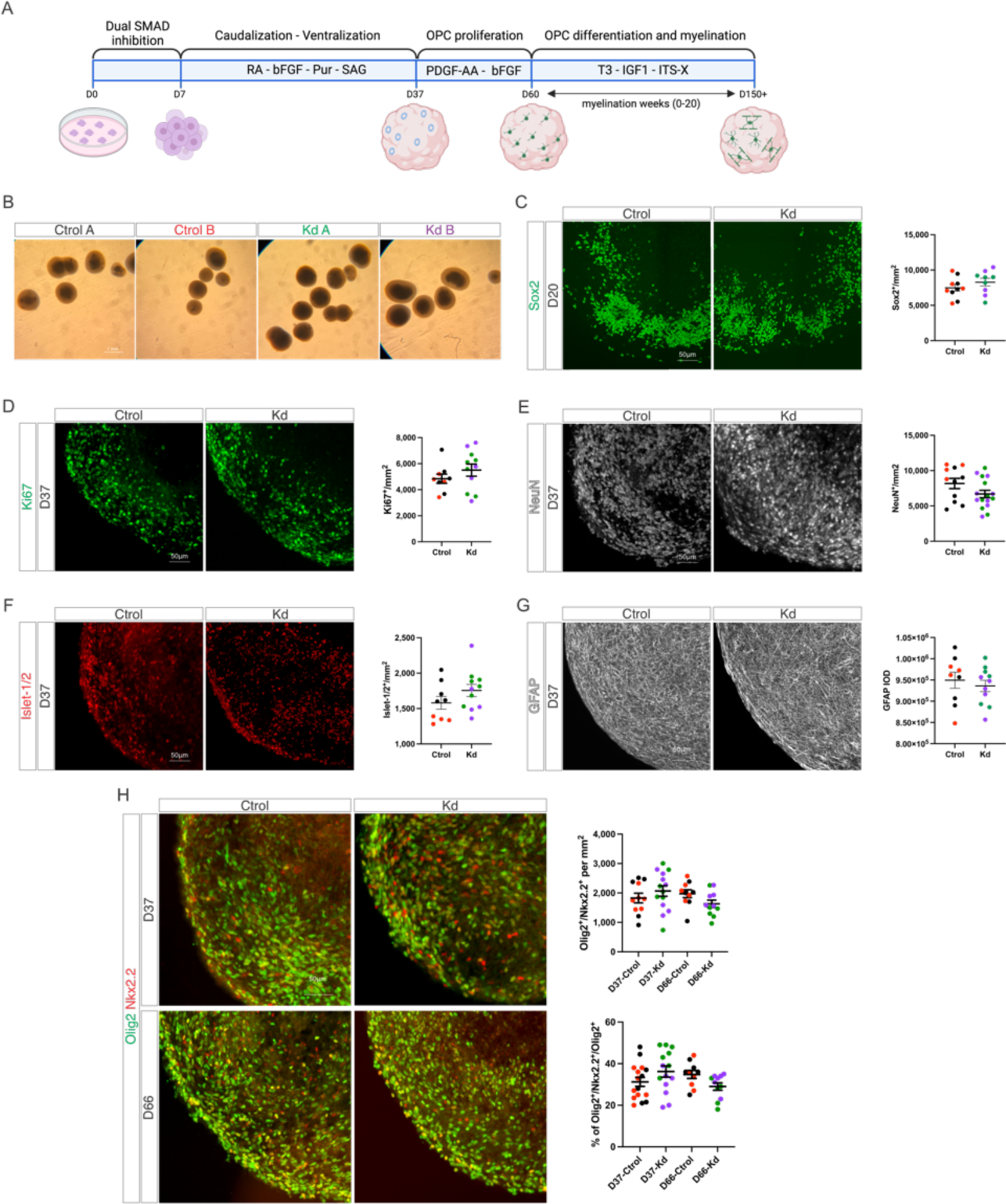
Kd organoids show normal neurogenesis, astrogenesis and oligodendrogenesis. (A) Graphical abstract of the protocol used to produce myelinating organoids. (B) Brightfield images of Ctrol and Kd organoids at D60 of culture. (C) Immunohistochemistry and quantifications of the number of Sox2^+^ neural stem cells in Ctrol and Kd organoids at D20. (D) Immunohistochemistry and quantifications of the number of Ki67^+^ proliferating progenitors in Ctrol and Kd organoids at D37. (E) Immunohistochemistry and quantifications of the number of NeuN^+^ pan-neuronal marker in Ctrol and Kd organoids at D37. (F) Immunohistochemistry and quantifications of the number of Islet1/2^+^ neuronal progenitors in Ctrol and Kd organoids at D37. (G) Immunohistochemistry and densitometric quantifications of GFAP^+^ astrocytes in Ctrol and Kd organoids at D37. (H) Immunohistochemistry and quantifications of the number of Olig2^+^/Nkx2.2^+^ OPCs in Ctrol and Kd organoids at D37 and D60. Values are expressed as the number of Olig2^+^/Nkx2.2^+^ per area (upper chart) or the number of Olig2^+^/Nkx2.2^+^ relative to the Olig2^+^ population (lower chart). Colors indicate replicates from individual iPSC lines. Values are expressed as the means ± SEMs.

### Kd organoids show signs of demyelination and minimal mTOR pathway perturbation

To analyze if GALC deficiency affects myelination extent, we cultured organoids in oligodendrocyte differentiation/myelinating media for up to 28 weeks after the oligodendrocyte proliferation phase (D60) and counted the number of MBP^+^ myelin internodes and internode’s length. Ctrol organoids show early signs of myelination after the 8^th^ week of exposure to myelination media and fully established myelin and Nodes of Ranvier after 12 weeks in this media (**Figure 3A**). After 12 weeks of myelination, Ctrol and Kd organoids show a similar number of MBP^+^ internodes, although a small decrease trend was observed in Kd organoids that show shorter internodes when compared to Ctrol ones (**Figure 3B**). Next, we cultured the organoids for another 8 weeks in myelination media (20 wk myelination) and found that Kd organoids have a decreased number of MBP^+^ internodes with persistent shorter internodes (**Figure 3B**). Interestingly, Ctrol organoids also show a trend decrease in the number of MBP^+^ internodes between weeks 12 and 20 of myelination, indicating that the culture conditions might have a limitation in maintaining myelin for long periods of time. To further explore this and to test if the decrease in the number of MBP^+^ internodes observed in Kd organoids becomes a stronger phenotype, we cultured Ctrol and Kd organoids for an additional 8 weeks in myelinating media (28 wk myelination) and found that both Ctrol and Kd organoids show extensive loss of myelin and a strong reduction in the number of Olig2^+^ oligodendrocytes (**Sup. Figure 1**). Ultrastructural EM analysis shows that Kd organoids have signs of lysosomal enlargement, vacuolation, lipid droplet (**Figure 3C** – upper panel) and myelin debris accumulation (**Figure 3C** – lower panel). Altogether, our findings show that the demyelinating phenotype observed in Kd can be replicated using iPSC-derived organoids *in vitro*.

**Figure 3.**
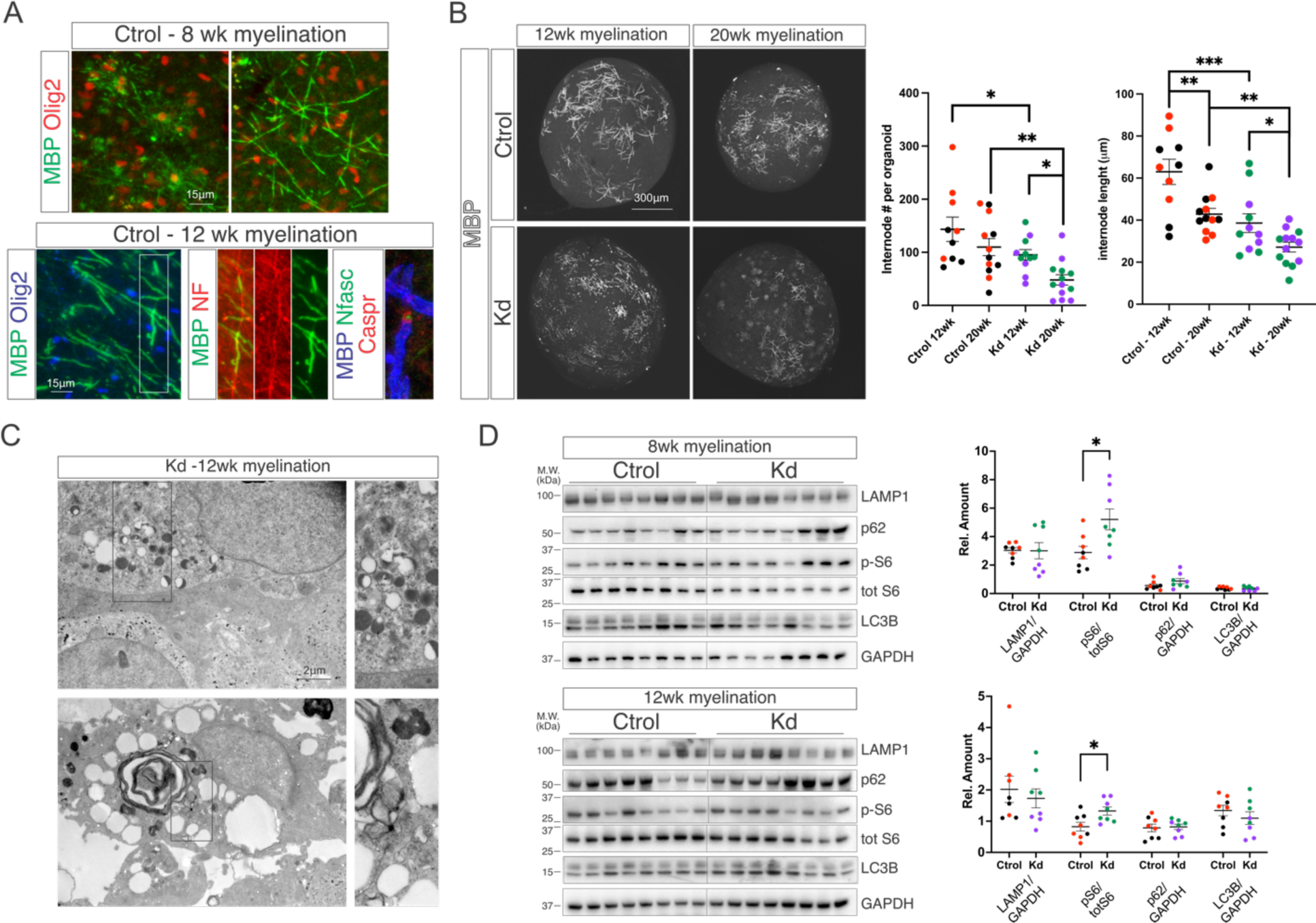
Kd organoids signs of demyelination and minimal mTOR pathway perturbation. (A) Immunohistochemistry for MBP^+^/Olig2^+^ mature oligodendrocytes at 8 or 12 weeks of myelination. Immunohistochemistry at 12 weeks for MBP/NF myelinated axons and MBP/Nfasc/Caspr fully developed nodes of Ranvier. (B) Immunohistochemistry for MBP in Ctrol and Kd organoids at 12 and 20 weeks of myelination. Quantifications of the number and length of MBP^+^ myelin internodes in Ctrol and Kd organoids at 12 and 20 weeks of myelination. (C) Electron micrographs of Kd organoids at 12 weeks of myelination showing lysosomal enlargement, vacuolation, lipid droplets and myelin debris. (D) Western blot of Ctrol and Kd organoid’s lysates at 8 and 12 weeks of myelination for the lysosomal marker LAMP1, the ribosomal protein S6 (S6) and the autophagy proteins p62 and LC3B. Colors indicate replicates from individual iPSC lines. Values are expressed as the means ± SEMs. \**P* < 0.05, \*\**P* < 0.01, and \*\*\**P* < 0.001 by Student’s *t* test or two-way ANOVA followed by Newman–Keuls multiple-comparisons post-test.

Kd is an LSD caused by the deficiency of GalC and as such, lysosomal dysfunction is expected to lead to defects in macromolecule’s recycling and metabolism. Our previous findings showed that an early myelination defect followed by myelin loss is present in Kd organoids starting at 12 weeks of myelination suggesting that perturbations in the above-mentioned processes might be present at 12 weeks of myelination or earlier. To evaluate this, we analyzed the content of the lysosomal marker LAMP1, the phosphorylation state of the ribosomal protein S6 (Inamura et al., 2018; Lin et al., 2023) and the autophagy markers p62 and LC3B (Del Grosso et al., 2019; Del Grosso et al., 2016; Papini et al., 2023). Western blot analysis of organoid lysates at 8 and 12 weeks of myelination show no differences in the content of LAMP1, p62 and LC3B, and a minor increase in the phosphorylation state of S6 at both time points (**Figure 3D**) indicating that the mentioned perturbations might be a late event in our model and not the cause for myelin loss.

At this point we were able to establish that Kd organoids lacking microglia/macrophages show an intrinsic, inflammation-independent, demyelinating phenotype that resembles the one observed in patients and animal models however, globoid cells are pathognomonic in Kd and because of this, we decided to co-culture our organoids with microglial progenitors. We obtained microglial progenitors by using a previously described protocol (Douvaras et al., 2017) (**Sup. Figure 2A**). After CD14 MACS we obtained pure Cx3CR1^+^/CD14^+^ microglia progenitors (D35) that differentiate into Iba1^+^ mature microglia after 15 days *in vitro* (D50) (**Sup. Figure 2B**). Next, we added Cx3CR1^+^/CD14^+^ microglial progenitors to D60 organoids, changed them into myelinating media and checked for the presence of microglia for an additional 3, 6 or 9 weeks (D60 + 3wk/6wk/9wk). At D60 + 3wk a significant number of Iba1^+^ microglia are present in organoids and show reactive morphology but at later time points, we observed a strong reduction in the number of Iba1^+^ microglia (**Sup. Figure 2C**) similar to what has been recently reported (Schafer et al., 2023), indicating a limitation of globoid cells *in vitro*.

### Globoid cell formation is drastically increased in Kd microglia by GalCer feeding

Globoid cells are prominently found in Kd tissues and suggested to be a key player in Kd’s progression (Favret et al., 2024). To test if the GALC deficient microglia can become globoid cells, we isolated Ctrol and Kd microglial progenitors, allowed them to differentiate into mature microglia, fed them with media containing GalCer or vehicle and counted the number of globoid-shaped cells labeled with Phalloidin. Kd microglia fed with vehicle show a minor but significant increase in the number globoid shaped cells (rounded and multinucleated cells) when compared to Ctrol ones however, a very drastic increase in this phenomenon is observed when Kd microglia are fed with GalCer for 24 or 48 hrs. (**Figure 4A**). A hallmark of globoid cells is their multinucleation, which is present in Kd cells after 24 hrs. of GalCer feeding and persisted after 48 hrs. of culture under these conditions (**Figure 4B**). Next, hypothesized that globoid cells are very likely to experience lysosomal dysfunction and this will manifest as defects in macromolecule’s recycling and cellular metabolism. To explore this idea, we analyzed the content of the lysosomal marker LAMP1, the phosphorylation state of the ribosomal protein S6 and the autophagy markers p62 and LC3B. First, we analyzed vehicle-fed Ctrol and Kd cells and found that the last ones have a slight decrease in LAMP1 content and a slight increase in LC3B content (**Figure 4C**). Next, we analyzed the effect of GalCer feeding at an early (16 hrs.) and a late time point (48 hrs.) and our findings show that Ctrol cells show minimal perturbation as reflected by a minor and transient increase in S6 phosphorylation state whereas Kd cells show drastic p62 and LC3B accumulation, reduction in LAMP1 content and a late increase in S6 phosphorylation (**Figure 4D**). All these suggest that initially, GalCer overload in Kd is detrimental to lysosomes but later on, globoid cells put in play compensatory mechanisms to increase the number of lysosomes and reduce the accumulation of phagocytic proteins.

**Figure 4.**
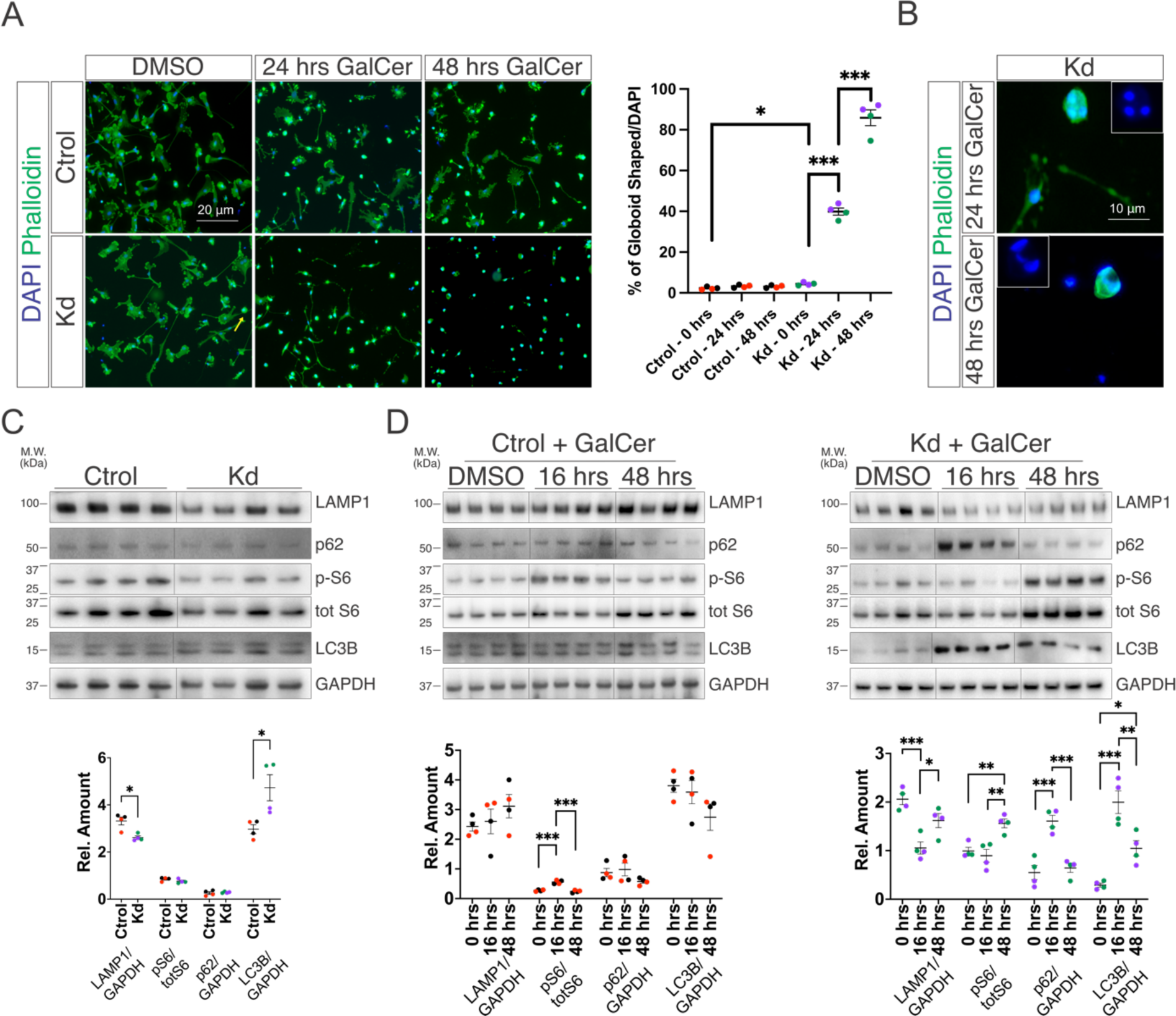
Globoid cell formation is drastically increased in Kd microglia by GalCer feeding. (A) Differentiated microglia (D50) from Ctrol and Kd iPSCs treated with vehicle (DMSO + HMCD) or 10 μM GalCer for 24 or 48 hrs. Phalloidins’ staining of actin to identify cell morphology. Quantification of the number of globoid shaped cells. Arrow indicates globoid cell in Kd cultures fed with DMSO. (B) Phalloidin staining for globoid multinucleated cells in Kd cultures exposed to GalCer for 24 or 48 hrs. (C) Western blot analysis of untreated Ctrol and Kd microglia for the lysosomal marker LAMP1, the ribosomal protein S6 (S6) and the autophagy proteins p62 and LC3B. (D) Western blot analysis of Ctrol (left) and Kd (right) microglia exposed to GalCer for 16 or 48 hrs. for the lysosomal marker LAMP1, the ribosomal protein S6 (S6) and the autophagy proteins p62 and LC3B. Colors indicate replicates from individual iPSC lines. Values are expressed as the means ± SEMs. \**P* < 0.05, \*\**P* < 0.01, and \*\*\**P* < 0.001 by Student’s *t* test.

## Discussion

Currently, Kd patients receive supportive care and the only approved disease-modifying treatment is hematopoietic stem cell transplantation (HSCT) which is effective in asymptomatic patients (Escolar et al., 2005; Krivit et al., 1998). Despite the immense success of HSCT in decreasing Kd’s progression, patients who receive HSCT experience variable levels of disability that are extensive in symptomatic patients (Wright et al., 2017; Yoon et al., 2021). In line with this, efforts have been made to explore additional treatments including substrate reduction therapy (SRT) (blocking catabolic pathways to prevent GalCer accumulation), enzyme replacement therapy (ERT) (administration of recombinant GalC) or gene therapy (viral delivery of GalC’s construct) (Heller et al., 2023). Among the single treatments, adeno-associated virus (AAV) delivery of GalC’s construct shows the best outcomes in disease progression and survival of pre-symptomatic mice and dogs (Hordeaux et al., 2022; Rafi et al., 2021). In addition, gene therapy combined with HSCT, shows superior disease outcomes in pre-symptomatic mice (Rafi et al., 2021; Rafi et al., 2015). Currently, there are two ongoing gene therapy-based clinical trials (NCT05739643 and NCT04693598) (https://clinicaltrials.gov/). In addition, combining three different treatments including HSCT, gene therapy and SRT given to pre-symptomatic mice showed even better disease outcomes than dual HSCT/gene therapy treatment (Li et al., 2021), but despite increasing the survival of the animals by 10-fold, motor dysfunction measured by rotarod and wire hanging tests, persists similarly to what happens in patients that received HSCT (Wright et al., 2017; Yoon et al., 2021) ultimately highlighting the importance of the need for a better understanding of the mechanisms that drive demyelination, neurodegeneration and neuroinflammation.

To identify the molecular mechanisms that underlay Kd pathology, iPSCs and iPSC-derived cultures were used. Liu and colleagues (Lv et al., 2023) performed bulk RNA sequencing of Kd iPSCs and NSC and found early dysregulated genes in Kd iPSCs and a significantly larger number of dysregulated genes in Kd NSC. Pathway analysis shows the involvement of novel pathways in Kd iPSCs, and the dysregulation of the MAPK, PI3K-Akt, cAMP, and RAS signaling pathways, as well as neuroactive ligand-receptor in Kd NSC. Interestingly, neither Kd iPSCs nor NSCs show dysregulation of the Kd associated Lysosome or Sphingolipid Metabolism KEGG pathways supporting the idea that the findings by Lv and colleagues are novel early events in Kd. Another work differentiated Kd iPSCs into astrocytes that show no gross development defects, accumulate psychosine, have increased expression and levels of IL-6, and a lipid accumulation phenotype (Lieberman et al., 2022). Interestingly, Kd astrocytes negatively impact the survival of Ctrol neurons (both in a co-culture setting or by adding astrocyte conditioned media) but positively impact Ctrol microglia survival in co-culture. Mangiameli and colleagues (Mangiameli et al., 2021) studied Kd patient-derived iPSCs and found that they retain normal stem cell properties, despite of accumulating psychosine, and show similar differentiation efficiency and comparable phenotype to Ctrol ones when differentiated into NSC. When Kd iPSCs were differentiated into neuronal/glial mixed cultures, the oligodendroglial and neuronal populations were negatively impacted. This phenomenon is explained by the activation of an early senescence program that affects lineage commitment, and an altered lipidomic profile throughout the iPSC to NSC stage. Altogether, these studies show that Kd impacts neurodevelopment in iPSC-derived 2D cultures but none of them explored myelination, which is largely impacted in Kd. In line with this, we used a 3D culture method that produces robust myelination (James et al., 2021) among the several approaches to induce iPSC differentiation into oligodendrocyte and myelination (Shim et al., 2023) and were able to replicate, for the first time, the demyelinating phenotype observed in Kd using iPSC-derived cultures.

Axonopathy in the spinal cord of pre-symptomatic Twitcher and Twitcher5J mice has been described before any detectable demyelination occurs suggesting a neuronal cell-autonomous phenotype (Castelvetri et al., 2011; Potter et al., 2013). Interestingly, these findings do not manifest as a reduction in the number of NeuN^+^ cells until the later stages of the disease. In support of a neuronal cell-autonomous mechanism of GALC ablation, early neuronal maturation defects together with a reduced number of Sox2^+^/Olig2^+^ OPCs have been shown to occur in the brain stem of GALC null mice (Weinstock, Kreher, et al., 2020) and a follow up study using pan-neuronal GALC deficient mice showed morphological signs of neurodegeneration in the spinal cord of 6 month old animals confirming a neuronal cell-autonomous phenotype in Kd (Kreher et al., 2022). Because of all of these, we can expect that our Kd organoids are very likely to experience some extent of axonopathy, that is not detectable by NeuN staining during the early stages of the cultures but could be detected at later stages, and to not show any neurogenesis defects because of recapitulating spinal cord development instead of brain stem’s. In this context, the results reported by Mangiameli and colleagues (Mangiameli et al., 2021) showing neurodevelopmental defects, but not found in our study, could be caused by the limitations of 2D as compared to 3D cultures (Jensen & Teng, 2020), the use of a protocol that yields a modest number of astrocytes and oligodendrocytes (limiting potential compensatory mechanisms arising from these cell populations) or because of recapitulating forebrain development whereas our protocol recapitulates spinal cord development, shows abundant number of both cell populations and replicates the findings described in mice.

In Kd, lysosomes fail to degrade GalCer resulting in the production and accumulation of psychosine which ultimately leads to lysosomal dysfunction (Folts et al., 2016). Lysosomes have multiple roles in several cellular processes including material recycling, metabolic signaling, autophagy, gene regulation, plasma membrane repair, and cell adhesion and migration (Ballabio & Bonifacino, 2020). In line with this, Kd phenotype has been linked to: metabolic dysregulation as shown by mTOR pathway dysregulation (Inamura et al., 2018; Lin et al., 2023) and autophagy impairment (Del Grosso et al., 2019; Del Grosso et al., 2016; Lin et al., 2023; Papini et al., 2023). Interestingly, our Kd organoids show signs of demyelination in the absence of any major autophagy dysregulation and the presence of a minor mTOR pathway dysregulation suggesting the absence of lysosomal dysfunction. Because of these, our Kd organoids are a valuable method to study the early events that drive demyelination in Kd in the absence of neuroinflammation, overcoming the limitations of using human biopsies. In regards to globoid cells. Kd microglia that weren’t fed with GalCer show small signs of what could be lysosomal impairment when compared to Ctrol microglia however, overloading microglia (both Ctrol and Kd) with GalCer shows an interesting phenomenon. In Ctrol microglia fed with GalCer, lysosomes can handle the overload as shown by the lack of accumulation of autophagic proteins and a modest mTOR pathway hyperactivation at an early time point. Later on, a trend decrease in autophagic proteins together with an increased lysosomal biogenesis, suggested by a trend increase of LAMP1 content, is observed and we interpreted this as normal compensation to lysosome’s overload. In the case of Kd microglia fed with GalCer, a strong accumulation of autophagic proteins and reduction of LAMP1 is observed at an early time point but at later on, when most cells had become globoid cells, a similar compensatory mechanism takes place by increasing LAMP1 content, reducing autophagic proteins and increasing mTOR pathway activation. Altogether this suggests that Kd lysosomes could be damaged at early time points, but replaced later on, showing that globoid cells possess the capacity to promote lysosome biogenesis in an attempt to restore autophagic flow which could be explored in the future using our model of globoid cells.

Globoid cells are pathognomonic of Kd, show a storage phenotype, are PAS^+^, have a distended cytoplasm with enlarged lysosomes, lipid crystals (presumably GalCer), and partially digested myelin sheaths (Nelson et al., 1963; Norman et al., 1961). In regards to how globoid cells form, macrophages/microglia fed with psychosine (Claycomb et al., 2014; Ijichi et al., 2013) or GalCer (Weinstock, Shin, et al., 2020) give rise to globoid cells. However, GalCer feeding induces globoid cells only in GALC deficient macrophages causing accumulation of both psychosine and GalCer whereas psychosine feeding induces globoid phenotype in wild type cell (wt) cultures. This indicates that psychosine is the driving force of globoid cell formation however, wt macrophages fed with GalCer (that do not become globoid cells) are very likely to resemble the foamy phagocytic cells observed in mice that received HSCT (Hoogerbrugge et al., 1988). These cells are a relevant cell population for treatment purposes like boosting the phagocytic capabilities of donor macrophages in LSDs (Weinstock, Shin, et al., 2020; Wolf et al., 2020). In this context, our model of globoid cells will allow us to study both globoid cells (Kd + GalCer) and overloaded donor-like cells (Ctrol + GalCer), in order to get a better understanding of the mechanisms that drive globoid cell formation and to identify novel mechanisms to promote donor cell’s beneficial effects.

## Acknowledgements

We would like to specially thank Dr. Daesung Shin for critical reading of the manuscript and all the members of our laboratory for constructive discussions. Also, we will like thank The Rosenau Family Foundation (formerly The Legacy of Angels Foundation) for their support.

## Figure legends

**Supplemental Figure 1.**
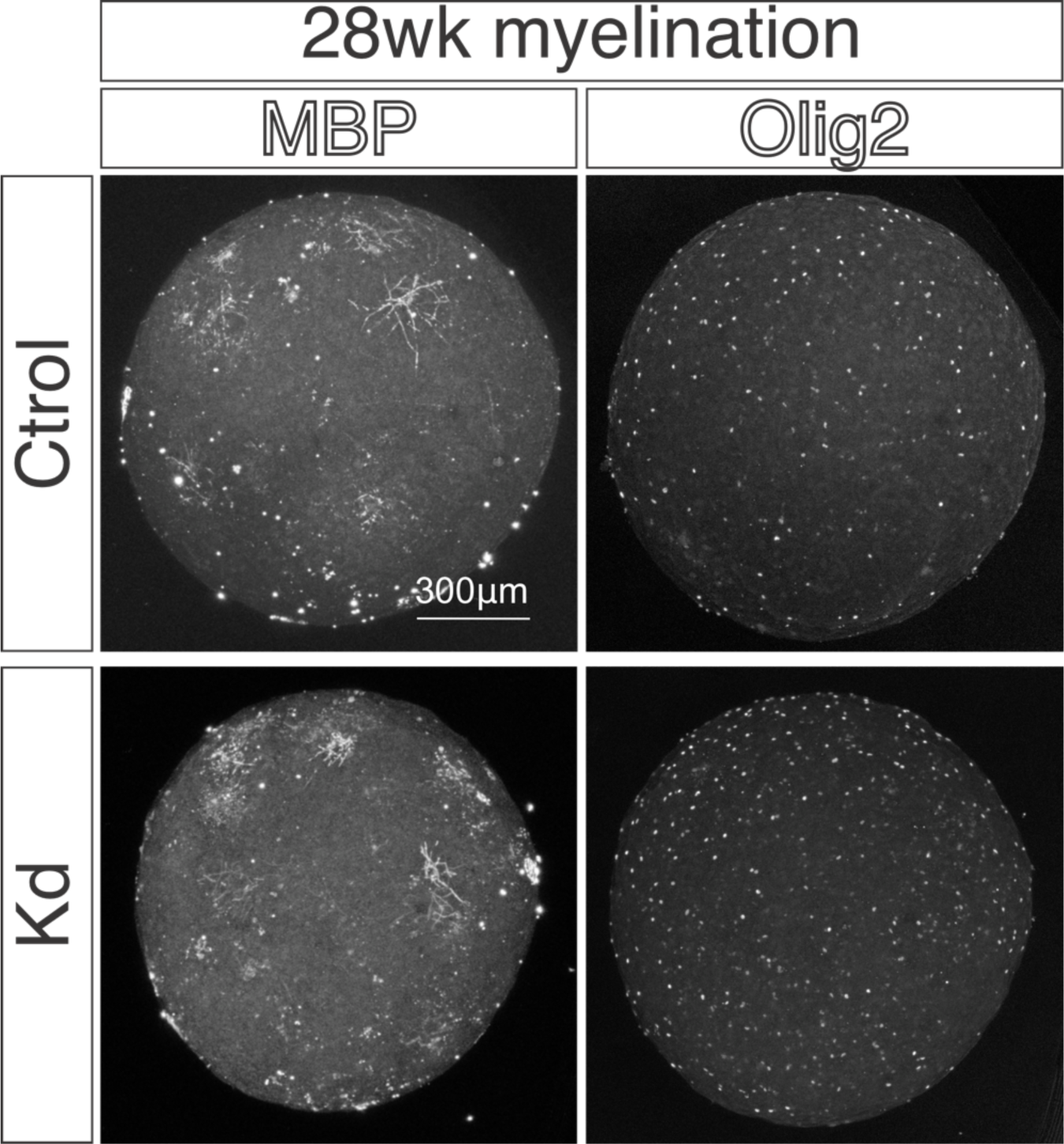
Both Ctrol and Kd organoids show strong myelin disruption at 28 weeks of myelination. Immunohistochemistry for MBP and Olig2 of Ctrol and Kd Organoids at 28 weeks of myelination.

**Supplemental Figure 2.**
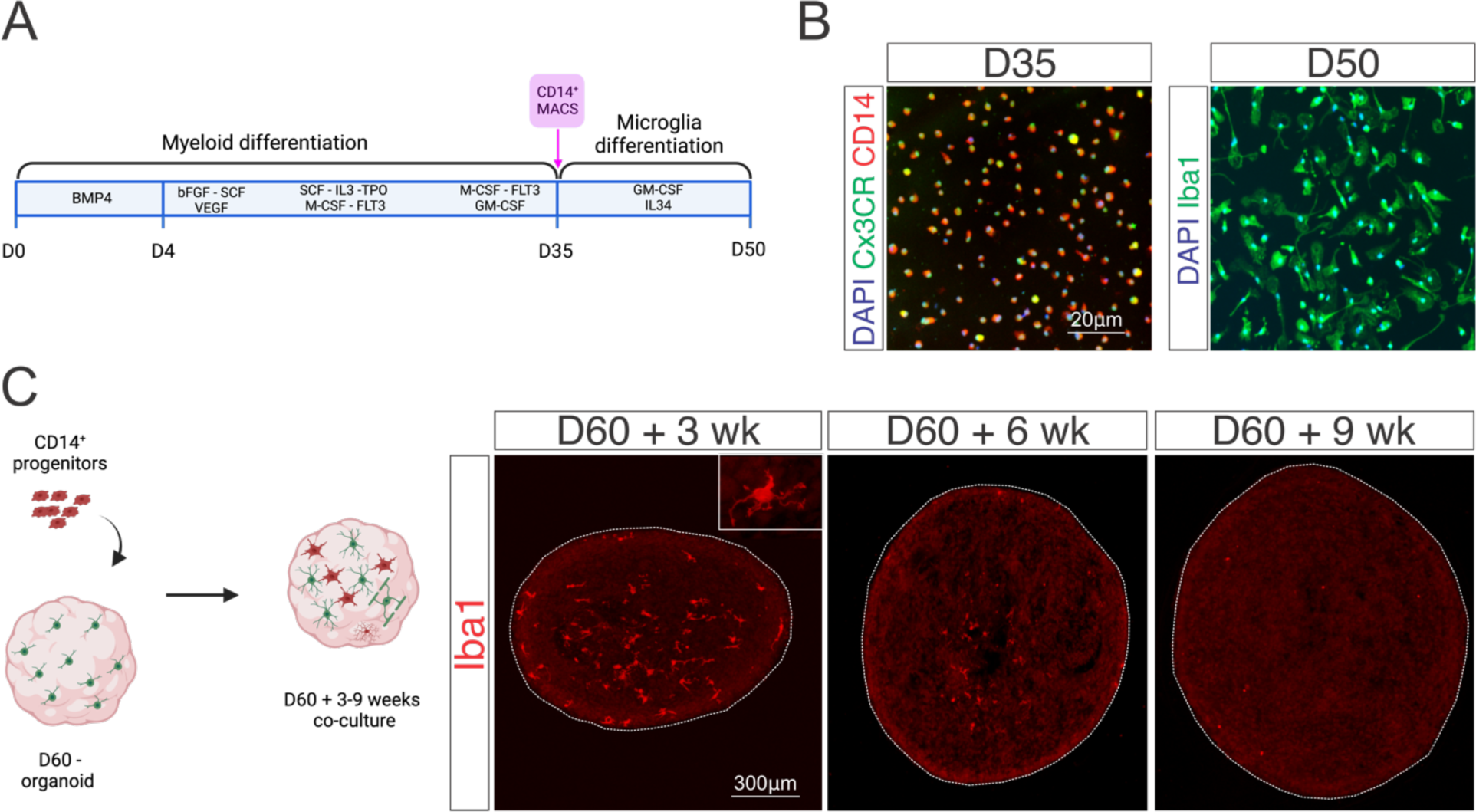
Microglial progenitors engraft organoids but do not survive beyond 6 weeks of co-culture. (A) Graphical abstract of the protocol used to produce iPSC derived microglia. (B) Immunohistochemistry for Cx3R1^+^/CD14^+^ microglial progenitors immediately after isolation (D35) and Iba1^+^ microglia after 14 days of differentiation (D50). (C) Microglial progenitors’ (D35) engraftment into D60 organoids. Immunohistochemistry for Iba1^+^ microglia engrafted into organoids 3, 6 and 9 weeks after engraftment.

